# Impaired intracellular Ca^2+^ signaling contributes to age-related cerebral small vessel disease in *Col4a1* mutant mice

**DOI:** 10.1101/2022.08.24.505186

**Authors:** Evan Yamasaki, Pratish Thakore, Sher Ali, Alfredo Sanchez Solano, Xiaowei Wang, Xiao Gao, Cassandre Labelle-Dumais, Myriam M. Chaumeil, Douglas B. Gould, Scott Earley

**Affiliations:** Department of Pharmacology, Center for Molecular and Cellular Signaling in the Cardiovascular System, University of Nevada, Reno School of Medicine, Reno, NV 89557-0318, USA; Department of Ophthalmology, UCSF School of Medicine, San Francisco, CA 94158, USA; Department of Physical Therapy and Rehabilitation Science, UCSF School of Medicine, San Francisco, CA 94143, USA; Department of Radiology and Biomedical Imaging, UCSF School of Medicine, San Francisco, CA 94143, USA; Department of Anatomy, Institute for Human Genetics, Cardiovascular Research Institute, Bakar Aging Research Institute, UCSF School of Medicine, San Francisco, CA 94158, USA

**Keywords:** *COL4A1*, collagen, cerebral small vessel diseases, basement membrane, myogenic tone, TRPM4, IP_3_R, Calcium, Ca^2+^, ER Stress, SR Stress, ion channels

## Abstract

Humans and mice with mutations in *COL4A1* and *COL4A2* manifest hallmarks of cerebral small vessel disease (cSVD), but the pathogenic mechanisms are largely unknown. Mice with a missense mutation in *Col4a1* at amino acid 1344 (*Col4a1^+/G1344D^*) exhibited age-dependent intracerebral hemorrhage (ICH) and brain lesions. Here we report that this pathology was associated with the loss of myogenic vasoconstriction, an intrinsic vascular response essential for the autoregulation of cerebral blood flow. Electrophysiological analyses showed that the loss of myogenic constriction resulted from blunted pressure-induced smooth muscle cell (SMC) membrane depolarization. Further, we found that dysregulation of membrane potential was associated with impaired Ca^2+^-dependent activation of large-conductance Ca^2+^-activated K^+^ (BK) and transient receptor potential melastatin 4 (TRPM4) cation channels linked to disruptions in sarcoplasmic reticulum (SR) Ca^2+^ signaling. Treating *Col4a1^+/G1344D^* mice with 4-phenylbutyrate, a compound that promotes the trafficking of misfolded proteins and alleviates SR stress, restored SR Ca^2+^ signaling, BK and TRPM4 channel activity, prevented loss of myogenic tone, and reduced ICH. We conclude that alterations in SR Ca^2+^ handling that impair membrane potential regulating ion channel activity result in dysregulation of SMC membrane potential and loss of myogenic tone contributing to age-related cSVD in *Col4a1^+/G1344D^* mice.

**Graphical Abstract:** 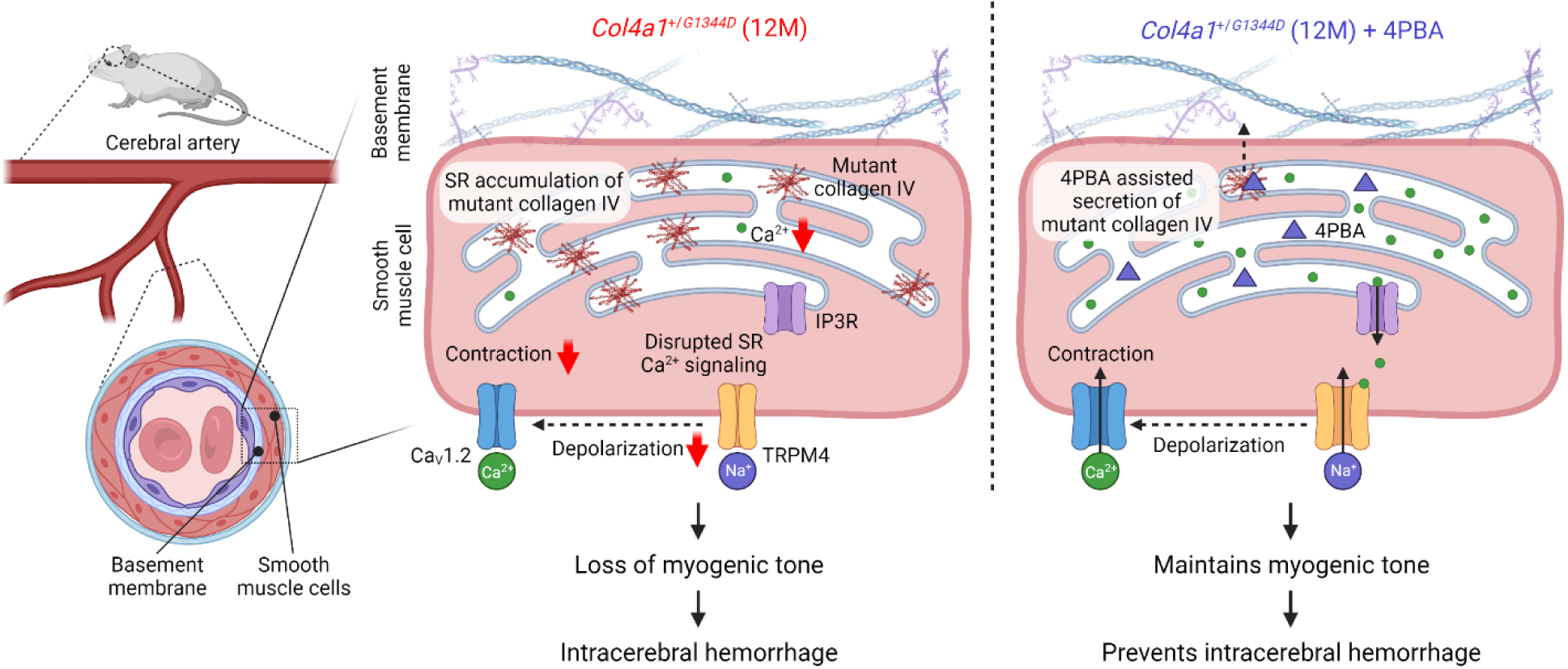

## Introduction

Cerebral small vessel diseases (cSVDs) are a group of related pathologies that damage arteries, arterioles, venules, and capillaries in the brain. cSVDs constitute a significant cause of stroke and, after Alzheimer’s disease, are the second leading cause of adult cognitive impairment and dementia ^1,2^. The underlying pathogenic mechanisms remain largely unknown, and no specific treatment options exist. Sporadic and familial forms of cSVD share clinical and radiological features, including white matter hyperintensities, dilated perivascular spaces, lacunar infarcts, microbleeds, and intracerebral hemorrhages (ICHs) ^3-6^. Individuals with Gould syndrome, a rare multisystem disorder caused by autosomal dominant mutations in collagen IV alpha 1 (*COL4A1*) and alpha 2 (*COL4A2*), manifest these hallmarks ^7-9^. The majority of pathogenic *COL4A1* and *COL4A2* mutations substitute critical glycine (G) residues within triple helical domains ^10^. These mutations have several deleterious effects, including impaired secretion of mature COL4A1 and COL4A2 heterotrimers [α1α1α2(IV)] that form basement membranes and intracellular retention of mutant collagen that can lead to endoplasmic reticulum (ER) stress ^11-13^. However, it is unclear how these defects influence the pathology of cSVD. *Col4a1* and *Col4a2* mutations demonstrate position-dependent effects contributing to phenotypic heterogeneity ^13^. For the current study, we investigated the pathogenesis of *Col4a1*-associated cSVD using mice with a missense mutation that changes the G residue at amino acid position 1344 to aspartate (D). These animals, designated *Col4a1^+/G1344D^*, model the most common type of mutation in humans and develop severe cSVD pathology compared to other mutant mice carrying glycine missense mutations in a murine allelic series of *Col4a1* and *Col4a2* mutations ^10,13^.

Distinctive attributes of the cerebral vasculature serve the unique requirements of the brain. For example, cerebral arteries and arterioles exhibit graded constriction in response to changes in intravascular pressure. This vital autoregulatory process, known as the vascular myogenic response, protects downstream capillary beds from potentially harmful increases in perfusion pressure when cardiac output is transiently elevated during exercise or other stimuli ^14^. The myogenic response is intrinsic to vascular smooth muscle cells (SMCs) and requires the coordinated activity of several ion channels. Increases in intraluminal pressure activate transient receptor potential melastatin 4 (TRPM4) channels, allowing an influx of Na^+^ ions that depolarizes the plasma membrane to increase voltage-dependent Ca^2+^ inflow through Cav1.2 Ca^2+^ channels and engage the contractile apparatus ^15,16^. Knockdown studies of TRPM4 show that *in vivo* suppression of TRPM4 decreases cerebral artery myogenic constriction and impairs cerebral blood flow autoregulation ^17^. Pressure-induced membrane depolarization is balanced by hyperpolarizing currents conducted by several K^+^-permeable ion channels, including large-conductance Ca^2+^-activated K^+^ (BK) channels, voltage-dependent K^+^ (K_V_) channels, and others ^18-20^. The interplay of depolarizing and hyperpolarizing currents controls the resting membrane potential of SMCs and maintains cerebral arteries in a state of partial contraction. BK and TRPM4 channels require high levels of intracellular Ca^2+^ for activation and, under native conditions, are stimulated by Ca^2+^ released from the sarcoplasmic reticulum (SR) through ryanodine receptors (RyRs) and inositol trisphosphate receptors (IP_3_Rs), respectively ^18,21^. Impairment of the myogenic response has been implicated in many types of cerebrovascular disease, including familial cSVDs associated with cerebral autosomal dominant arteriopathy with subcortical infarcts and leukoencephalopathy (CADASIL) caused by mutations in *NOTCH3* ^22^. However, the impact of the *Col4a1^G1344D^* mutation on the myogenic response is not known.

Here, we report that *Col4a1^+/G1344D^* mice exhibit age-dependent loss of myogenic tone that is associated with spontaneous intracerebral hemorrhage (ICH) and brain lesions detected by susceptibility-weighted magnetic resonance imaging (SWI). The deficit in myogenic tone resulted from diminished stretch-induced activation of TRPM4 channels and subsequent impairment of SMC membrane depolarization. We also found that subcellular and global Ca^2+^ signals generated by releasing Ca^2+^ from the SR through RyRs and IP_3_Rs were disrupted in SMCs from aged *Col4a1^+/G1344D^* mice. Treating *Col4a1^+/G1344D^* mice with 4-phenylbutyrate (4PBA), a small molecule with chaperone properties that improves the secretion of misfolded proteins from the ER/SR ^10,13,23-26^, prevented SR Ca^2+^ signaling defects, preserved TRPM4 channel activity, attenuated the loss of myogenic tone, and largely protected mice from ICH. These data provide evidence that accumulation of misfolded α1α1α2(IV) collagen in the SR of SMCs from *Col4a1^+/G1344D^* mice causes disruptions in intracellular Ca^2+^ signaling, stretch-induced activation of TRPM4 channels, pressure-induced SMC membrane potential depolarization, and myogenic constriction. Our data also suggests these defects contribute to spontaneous age-related ICH in *Col4a1^+/G1344D^* mice. Thus, the findings of this study reveal a novel age-dependent facet of cSVD pathogenesis.

## Results

### *Col4a1^+/G1344D^* mice exhibit age-dependent pathology

All studies utilized *Col4a1^+/G1344D^* mice and *Col4a1^+/+^* littermates as controls. *Col4a1^+/G1344D^* mice carry missense mutation at amino acid 1344 near the C-terminus of the triple helical domain (Figure 1A). Age is the most critical risk factor for cSVD and vascular dementia ^27,28^. To investigate age-related aspects of cSVD pathology, mice were studied at 3 and at least 12 months (M) of age, representing the young adult and middle age stages of life, respectively. Male and female mice were used throughout the study, and no sex-specific differences were detected.

**Figure 1.**
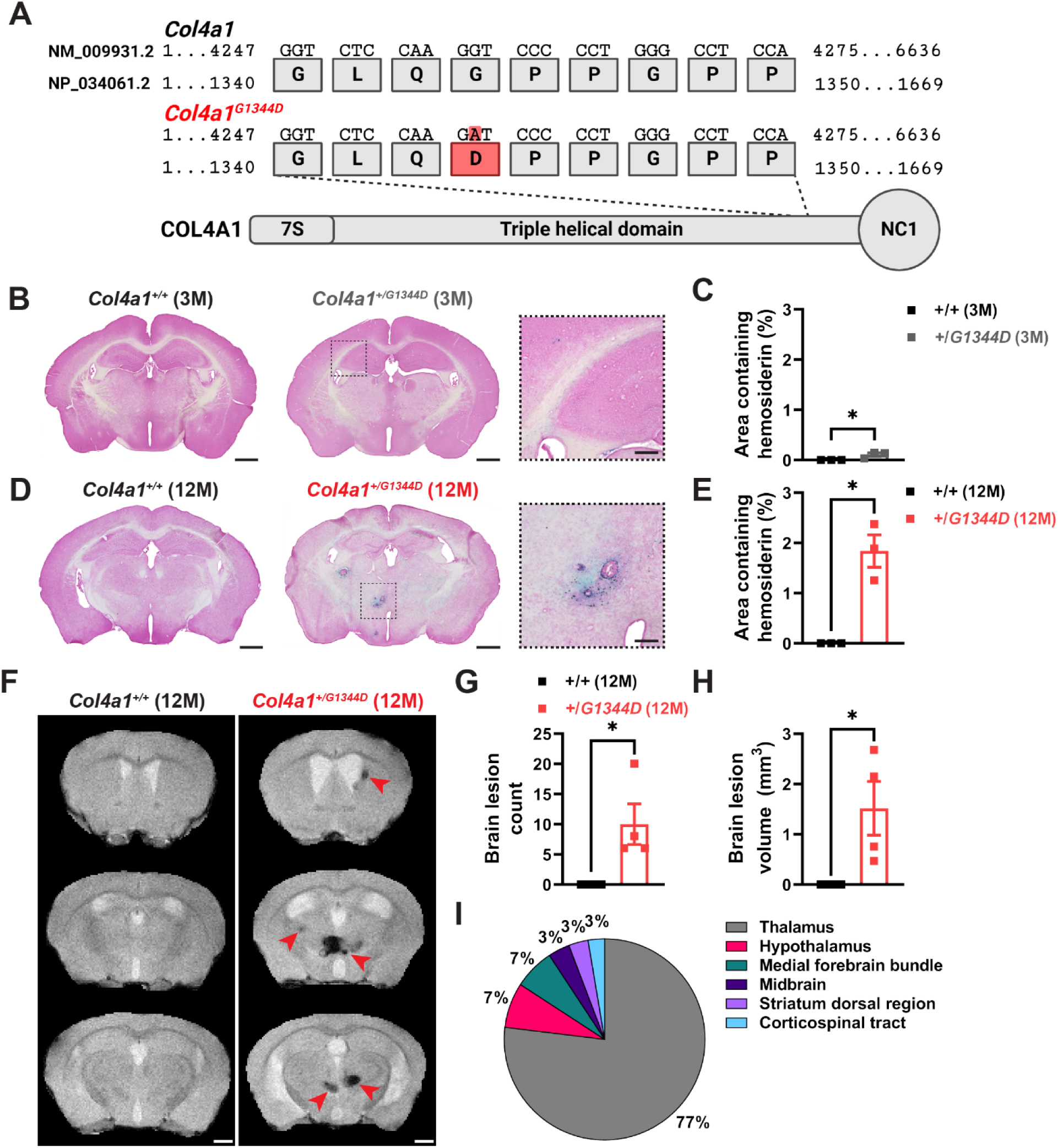
*Col4a1^+/G1344D^* mice exhibit age-dependent pathologies. **(A)** Schematic representation of the *Col4a1* point mutation that leads to *Col4a1^G1344D^*. **(B)** Intracerebral hemorrhage (ICH) was assessed using Prussian blue staining. Representative images of brain sections from 3 M-old *Col4a1^+/+^* and *Col4a1^+/G1344D^* mice stained with Prussian blue. **(C)** Summary data presented as percentage of brain area with Prussian blue staining in brain sections (500 µm intervals). N = 3 animals per group. *P ≤ 0.05, unpaired t-test. **(D)** Representative images of brain sections from 12 M-old *Col4a1^+/+^* and *Col4a1^+/G1344D^* mice stained with Prussian blue. **(E)** Summary data presented as percentage of brain area with Prussian blue staining in brain sections (500 µm intervals). N = 3 animals per group. *P ≤ 0.05, unpaired t-test. **(F)** Brain lesions were evaluated *in vivo* using high-field susceptibility-weighted magnetic resonance imaging (SWI). Representative SWI images from 12 M-old *Col4a1^+/+^* and *Col4a1^+/G1344D^* mice. Red arrowheads highlight hypointense pixels that indicate hemorrhagic lesions. Quantification of **(G)** the number of lesions and **(H)** the total volume of lesions detected by SWI. N = 4 animals per group. *P ≤ 0.05, unpaired t-test. **(I)** Region-wise volumetric distribution of lesions detected by SWI from 12 M-old *Col4a1^+/G1344D^* mice. Scale bars = 1 mm. Inset scale bars = 250 µm.

Spontaneous ICH is a highly penetrant and consequential manifestation of *Col4a1* mutations that increases in severity with age ^10,29,30^. ICH severity was measured in brain sections from *Col4a1^+/+^* and *Col4a1^+/G1344D^* mice stained with Prussian blue. ICH was not detected in 3 M-old *Col4a1^+/+^* mice. Low-level, punctate Prussian blue staining was present in brain sections from 3 M-old *Col4a1^+/G1344D^* mice (Figure 1B and C). 12 M-old *Col4a1^+/+^* mice did not exhibit ICH, but brain sections from 12 M-old *Col4a1^+/G1344D^* mice displayed large areas of Prussian blue staining mainly localized to the subcortical regions (Figure 1D and E). The total area of Prussian blue staining in 12 M-old *Col4a1^+/G1344D^* mice was nearly 20-fold greater than in 3 M-old mutants (1.9% vs. 0.1% of brain area, respectively), indicating that the severity of spontaneous ICH increases dramatically with age.

Neuroimaging is frequently used to diagnose cSVD in human patients ^31^. In this study, 12 M-old *Col4a1^+/+^* and *Col4a1^+/G1344D^* mice were evaluated using SWI. 12 M-old *Col4a1^+/+^* mice did not exhibit SWI-detectable brain lesions, but brains from age-matched *Col4a1^+/G1344D^* mice had multiple lesions, mainly localized to the thalamus (Figure 1F-I). The number, total volume, and localization of brain lesions detected with SWI are consistent with ICH detected with Prussian blue staining.

### Diminished pressure-induced constriction of cerebral arteries from 12 M-old ***Col4a1^+/G1344D^* mice is due to impaired SMC membrane depolarization.**

To better understand the underlying causes of ICH in *Col4a1^+/G1344D^* mice, we used *ex vivo* pressure myography to study the myogenic response of cerebral arteries ^32^. Vasoconstriction in response to stepwise increases in intraluminal pressure (from 5 to 120 mmHg) was evaluated by measuring the active steady-state luminal diameter of cannulated cerebral arteries bathed in standard physiological saline solution at each pressure. This procedure was then repeated using a Ca^2+^-free bathing solution to determine the passive response to pressure, and myogenic tone was calculated as the difference in active versus passive diameter normalized to the passive diameter. The myogenic tone of cerebral arteries isolated from 3 M-old *Col4a1^+/+^* and *Col4a1^+/G1344D^* mice did not differ (Figure 2A and B). However, cerebral arteries from 12 M-old *Col4a1^+/G1344D^* mice barely constricted in response to pressure and had significantly lower levels of myogenic tone than those from age-matched *Col4a1^+/+^* mice (Figure 2C and D).

**Figure 2.**
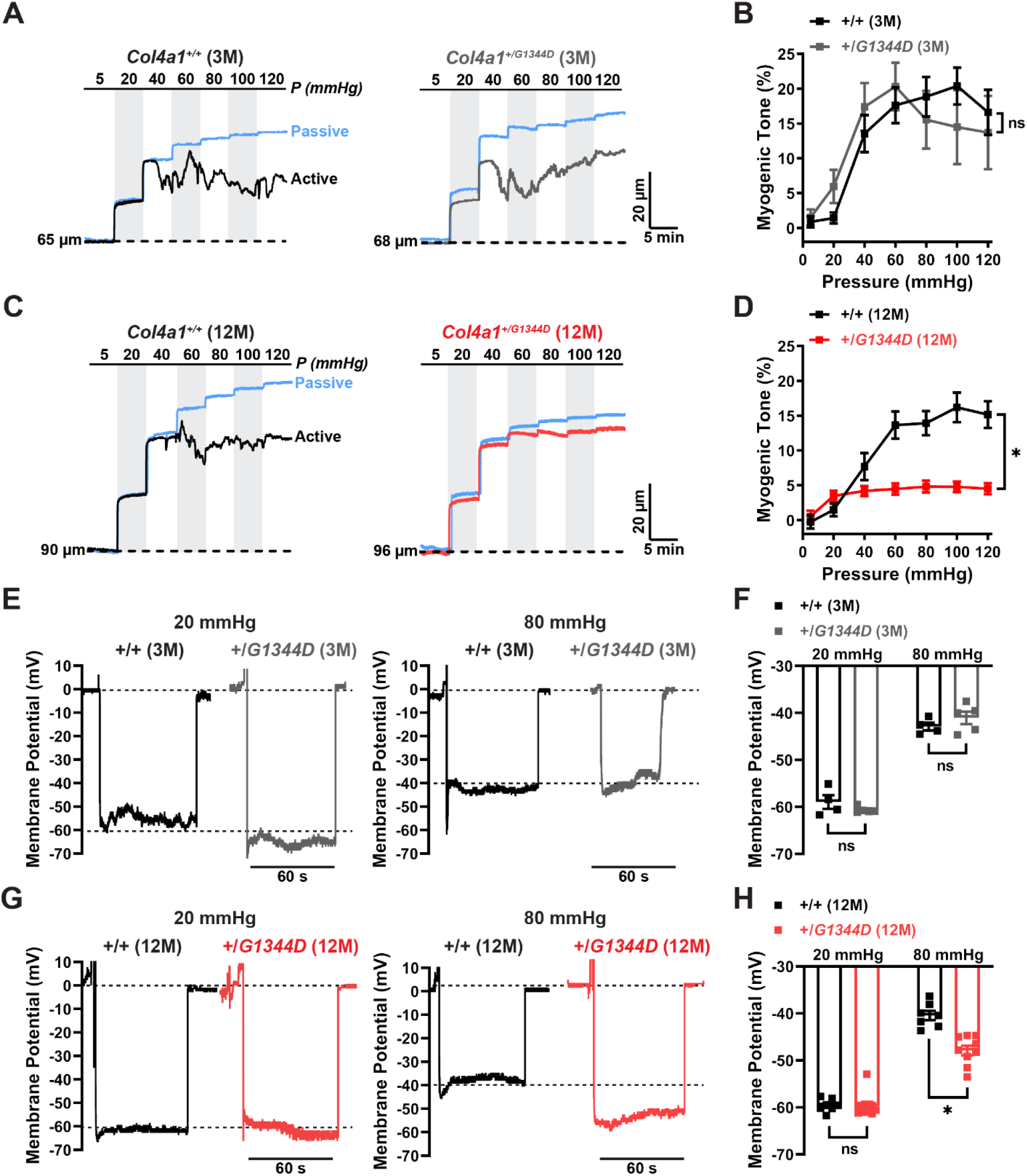
Diminished pressure-induced constriction of cerebral arteries from 12 M-old *Col4a1^+/G1344D^* mice is due to impaired SMC membrane depolarization. **(A)** Representative recordings of the inner diameter of isolated cerebral arteries from 3 M- old *Col4a1^+/+^* and *Col4a1^+/G1344D^* mice show the myogenic response to increased pressure (active) and the dilation of the same arteries when extracellular Ca^2+^ has been removed (passive). **(B)** Summary data of myogenic tone as a function of intraluminal pressure. n = 5-10 vessels from 5 or 6 animals per group. ns = not significant, two-way ANOVA. **(C)** Representative recordings of the inner diameter of isolated cerebral arteries from 12 M-old *Col4a1^+/+^* and *Col4a1^+/G1344D^* mice. **(D)** Summary data of myogenic tone as a function of intraluminal pressure. n = 13 vessels from 6 or 9 animals per group. *P ≤ 0.05, two-way ANOVA. **(E)** Representative membrane potential (mV) recordings of smooth muscle cells (SMCs) in pressurized cerebral arteries isolated from 3 M-old *Col4a1^+/+^* and *Col4a1^+/G1344D^* mice at 20 mmHg and 80 mmHg intraluminal pressure. **(F)** Summary data showing membrane potential of SMCs in pressurized cerebral arteries at 20 mmHg and 80 mmHg intraluminal pressure. n = 4-5 vessels from 3 or 4 animals per group. ns = not significant, unpaired t-test. **(G)** Representative membrane potential recordings of SMCs in pressurized cerebral arteries isolated from 12 M-old *Col4a1^+/+^* and *Col4a1^+/G1344D^* mice at 20 mmHg and 80 mmHg intraluminal pressure. **(H)** Summary data showing membrane potential of SMCs in pressurized cerebral arteries at 20 mmHg and 80 mmHg intraluminal pressure. n = 7-9 vessels from 5 or 6 mice per group. *P ≤ 0.05, unpaired t-test.

Increases in intraluminal pressure cause SMCs in the vascular wall to depolarize ^33^, triggering voltage-dependent Ca^2+^ influx and muscular contraction ^33,34^. To determine if the loss of myogenic tone in 12 M-old *Col4a1^+/G1344D^* mice was associated with impaired pressure-induced depolarization, we recorded the membrane potentials of SMCs in intact cerebral arteries using intracellular microelectrodes. At low intraluminal pressure (20 mmHg), the membrane potential of SMCs in cerebral arteries from 3 M-old *Col4a1^+/+^* and *Col4a1^+/G1344D^* mice did not differ (Figure 2E and F). Increasing intraluminal pressure to physiological levels (80 mmHg) depolarized the SMC membrane potential in arteries from both groups to the same extent (Figure 2E and F). When vessels from 12 M-old animals were investigated, SMC membrane potential did not differ between *Col4a1^+/+^* and *Col4a1^+/G1344D^* when arteries were pressurized to 20 mmHg (Figure 2G and H), but when intraluminal pressure was increased to 80 mmHg, the depolarization of SMCs in cerebral arteries from *Col4a1^+/G1344D^* mice was significantly blunted compared with controls (Figure 2G and H). These data suggest that cerebral arteries from 12 M-old *Col4a1^+/G1344D^* mice lose myogenic tone because pressure-induced SMC membrane depolarization is impaired.

### Age-dependent declines in BK and TRPM4 channel activity in SMCs from ***Col4a1^+/G1344D^* mice.**

To determine if changes in ion channel activity account for impaired pressure-induced SMC depolarization, we used patch-clamp electrophysiology to compare hyperpolarizing K_V_ and BK currents and depolarizing TRPM4 currents in native SMCs from control and *Col4a1^+/G1344D^* mice.

Elevated SMC K_V_ channel current density is responsible for diminished myogenic tone in cerebral arteries from CADASIL cSVD mice ^22^. We measured K_V_ currents in freshly isolated SMCs to determine if a similar mechanism accounts for the loss of myogenic tone in cerebral arteries from 12 M-old *Col4a1^+/G1344D^* mice. K_V_ currents were recorded using the amphotericin B perforated patch-clamp configuration to maintain the integrity of intracellular signaling cascades. Whole-cell currents were evoked by applying voltage steps from −60 to +60 mV from a holding potential of −80 mV in the presence of selective BK channel blocker paxilline (1 µM). The voltage step protocol was repeated in the presence of K_V_ channel blocker 4-aminopyridine (5 mM), and K_V_ current amplitude was determined at each potential by subtraction. We found that K_V_ current amplitudes did not differ between SMCs from 12 M-old *Col4a1^+/+^* and *Col4a1^+/G1344D^* mice at all applied command potentials (Supplemental Figure 1A and B).

The activity of BK channels in cerebral artery SMCs under native conditions is driven by the transient release of Ca^2+^ ions through RyRs on the SR into restricted subcellular microdomains proximal to the plasma membrane ^35^. These signaling events, known as Ca^2+^ sparks, activate clusters of BK channels on the plasma membrane to generate large-amplitude spontaneous transient outward currents (STOCs) ^18,19^. We utilized the amphotericin B perforated patch-clamp configuration to measure STOCs in cerebral artery SMCs from 3 M-old *Col4a1^+/+^* and *Col4a1^+/G1344D^* mice over a range of command potentials (−60 to 0 mV) and saw no significant difference in STOC frequency or amplitude (Figure 3A-C). However, the frequency and amplitude of STOCs were significantly lower in cerebral artery SMCs from 12 M-old *Col4a1^+/G1344D^* mice compared with age-matched controls (Figure 3D-F). To determine if decreases in STOCs result from changes in gene expression, we utilized ddPCR to measure transcript levels of *Kcnma1* (BK subunit α), *Kcnmb1 (*BK subunit β1), and *Ryr2* (ryanodine receptor 2). No differences in mRNA expression levels were detected (Supplemental Figure 2).

**Figure 3.**
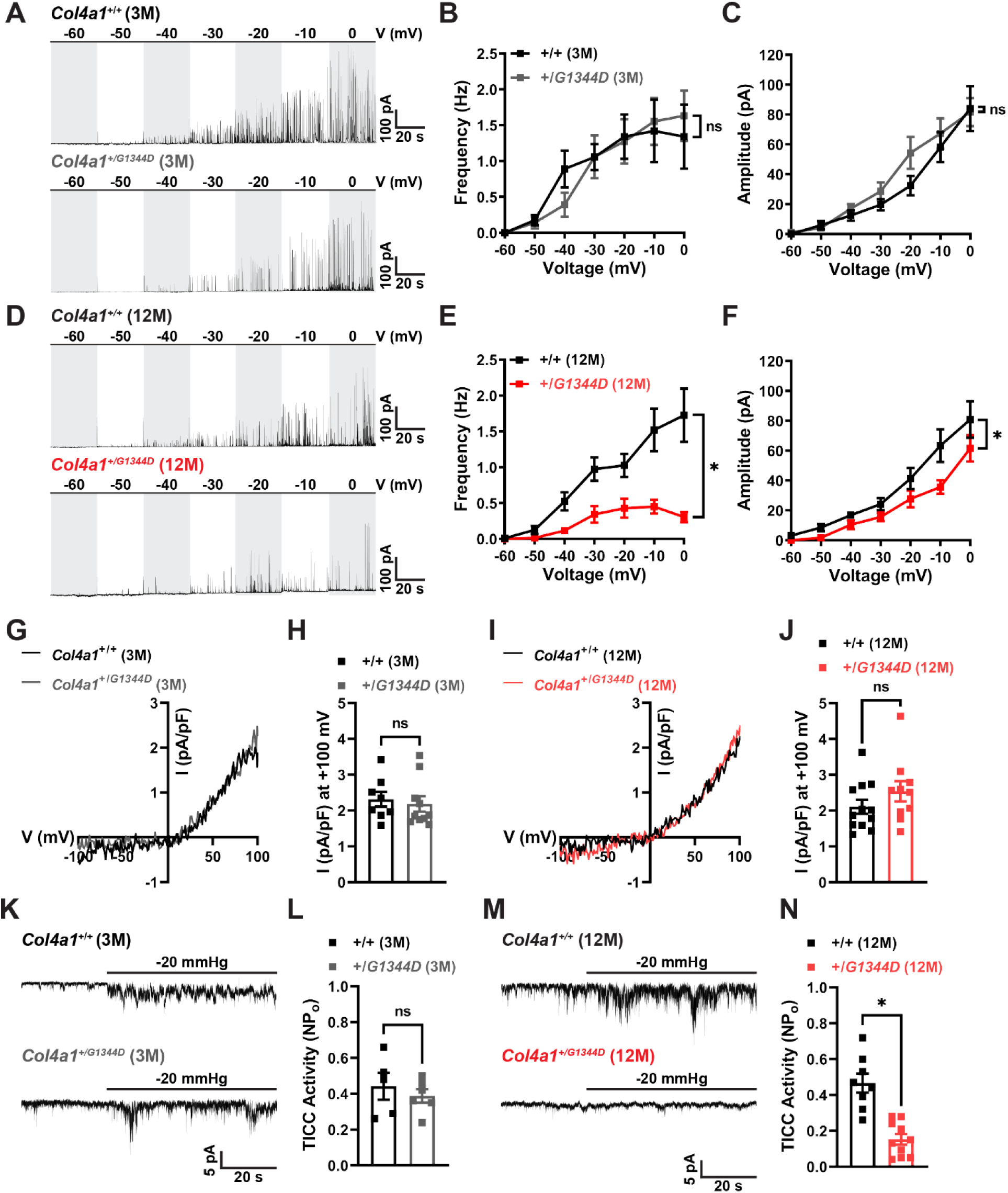
Age-dependent declines in BK and TRPM4 channel activity in SMCs from *Col4a1^+/G1344D^* mice. **(A)** Representative traces of spontaneous transient outward currents (STOCs) in freshly isolated cerebral artery smooth muscle cells (SMCs) from 3 M-old *Col4a1^+/+^* and *Col4a1^+/G1344D^* mice over a range of membrane potentials (−60 to 0 mV). Summary data showing STOC frequency **(B)** and amplitude **(C)** at each command potential. n = 8-12 cells from 5 animals per group. ns = not significant, two-way ANOVA. **(D)** Representative traces of STOCs in cerebral artery SMCs from 12 M-old *Col4a1^+/+^* and *Col4a1^+/G1344D^* mice. Summary data showing STOC frequency **(E)** and amplitude **(F)** at each command potential. n = 12 cells from 5 animals per group. *P ≤ 0.05, two-way ANOVA. **(G)** Representative I-V plots of whole-cell patch-clamp TRPM4 current recordings during voltage ramps (−100 to 100 mV) in cerebral artery SMCs from 3 M-old *Col4a1^+/+^* and *Col4a1^+/G1344D^* mice. Currents were evoked by including 200 µM free Ca^2+^ in the intracellular solution, and the TRPM4 portion of the current was determined by adding TRPM4 blocker 9-phenanthrol (30 µM). **(H)** Summary of whole-cell TRPM4 current density at +100 mV. n = 8-10 cells from 3 or 4 animals per group. ns = not significant, unpaired t-test. **(I)** Representative I-V plots of whole-cell patch-clamp TRPM4 current recordings in cerebral artery SMCs from 12 M-old *Col4a1^+/+^* and *Col4a1^+/G1344D^* mice. **(J)** Summary of whole-cell TRPM4 current density at +100 mV. n = 10-12 cells from 4 animals per group. ns = not significant, unpaired t-test. **(K)** Representative traces of transient inward cation currents (TICCs) evoked by applying negative pressure (−20 mmHg) to stretch the plasma membrane of voltage-clamped (−70 mV) cerebral artery SMCs isolated from 3 M-old *Col4a1^+/+^* and *Col4a1^+/G1344D^* mice. **(L)** Summary data of TICC activity. n = 5-6 cells from 3 animals per group. ns = not significant, unpaired t-test. **(M)** Representative traces of TICCs in cerebral artery SMCs isolated from 12 M-old *Col4a1^+/+^* and *Col4a1^+/G1344D^* mice. **(N)** Summary data of TICC activity. n = 8-10 cells from 4 or 5 animals per group. *P ≤ 0.05, unpaired t-test.

Reduced BK channel activity depolarizes the SMC membrane potential and increases contractility ^36^. Thus, diminished BK channel activity in SMCs from 12 M-old *Col4a1^+/G1344D^* mice cannot account for impaired pressure-induced membrane depolarization.

TRPM4 is a Ca^2+^-activated, nonselective monovalent cation channel required for pressure-induced SMC depolarization and the development of myogenic tone in cerebral arteries ^15,16,37,38^. At the membrane potentials of SMCs in the vascular wall under physiological conditions (−60 to −30 mV), TRPM4 channels conduct inward Na^+^ currents that depolarize the plasma membrane in response to increases in intraluminal pressure ^15,39^. Patch-clamp electrophysiology was used to determine if TRPM4 activity is diminished in *Col4a1^+/G1344D^* mice. We assessed channel function and availability in freshly isolated cerebral artery SMCs using the conventional whole-cell patch-clamp configuration with a high [Ca^2+^] intracellular solution to directly activate TRPM4. Currents were recorded as voltage ramps (−100 to +100 mV) were applied. Voltage ramps were repeated in the presence of TRPM4 blocker 9-phenanthrol (30 µM) ^15^, and TRPM4 currents were isolated by subtraction. Prior studies using ion substitution protocols have shown that Ca^2+^-activated, 9-phenanthrol currents recorded using these methods are carried by Na^+^ ions, providing substantial evidence that these are *bona fide* TRPM4 currents ^16,37,40^. We found that the amplitudes of whole-cell TRPM4 currents recorded from cerebral artery SMCs did not differ between *Col4a1^+/+^* and *Col4a1^+/G1344D^* mice at 3 and 12 M of age (Figure 3G-J). These data indicate that the collagen mutation studied here has no direct effects on TRPM4 channel function or the trafficking of channel protein to the plasma membrane.

TRPM4 channels in SMCs are activated by a signal transduction cascade initiated by angiotensin II type 1 receptors (AT_1_Rs) ^41-44^. Activation of Gq protein-coupled AT_1_Rs stimulates phospholipase C (PLC), leading to the cleavage of phosphatidylinositol 4,5-bisphosphate (PIP_2_) into diacylglycerol and inositol triphosphate (IP_3_). IP_3_ stimulates Ca^2+^ release from the SR through IP_3_Rs to transiently activate TRPM4 channels ^21,38,41^. AT_1_Rs are directly activated by the stretch of the plasma membrane (independent of their ligand) and are a critical mechanosensor in SMCs ^41-44^. To measure the activity of TRPM4 channels in response to mechanical force, we patch-clamped SMCs in the amphotericin B perforated patch configuration and applied negative pressure (−20 mmHg) through the patch pipet to stretch the plasma membrane. Stretch-induced transient inward cation currents (TICCs) evoked in this manner are blocked by 9-phenanthrol and abolished by the knockdown of TRPM4 expression ^15,21^. Stretch-activated TICC activity in SMCs from 3 M-old *Col4a1^+/+^* and *Col4a1^+/G1344D^* (Figure 3K and L) did not differ. In contrast, stretch-induced TICC activity in SMCs from 12 M-old *Col4a1^+/G1344D^* mice was significantly lower than that of age-matched *Col4a1^+/+^* mice (Figure 3M and N). To determine if altered gene expression underlies the reduction in TICC activity, we utilized ddPCR to measure transcript levels of *Trpm4* (TRPM4)*, Itpr1* (IP_3_ receptor 1), and *Itpr2* (IP_3_ receptor 2) in cerebral arteries from 12 M- old *Col4a1^+/+^* and *Col4a1^+/G1344D^* mice. No differences in mRNA copy numbers were observed (Supplemental Figure 2). These data suggest that blunted pressure-induced SMC depolarization and impaired myogenic tone result from diminished stretch-induced activation of TRPM4 channels.

### Defective intracellular Ca^2+^ signaling in cerebral artery SMCs from 12 M-old ***Col4a1^+/G1344D^* mice.**

Ca^2+^ released from the SR drives BK and TRPM4 channel activity in cerebral artery SMCs ^18,21^. Therefore, we investigated the possibility that these vital Ca^2+^ signaling pathways are disrupted in SMCs from 12 M-old *Col4a1^+/G1344D^* mice. Freshly isolated SMCs were loaded with the Ca^2+^-sensitive fluorophore Fluo-4 AM and imaged using high-speed, high-resolution spinning disk confocal microscopy. Spontaneous Ca^2+^ sparks were present in SMCs isolated from 3 M-old *Col4a1^+/+^* and *Col4a1^+/G1344D^* mice. The frequency, amplitude, duration, rise time, and decay rate did not differ between groups (Figure 4A-C). Spontaneous Ca^2+^ sparks were also present in SMCs isolated from both groups of 12 M-old mice. No significant differences in Ca^2+^ spark amplitude, duration, rise time, and decay rate were observed, but notably, Ca^2+^ spark frequency was significantly lower in SMCs from 12 M-old *Col4a1^+/G1344D^* mice (Figure 4D-F). Thus, diminished Ca^2+^ spark frequency likely accounts for decreased STOC frequency in SMCs from 12 M-old *Col4a1* mutant mice.

**Figure 4.**
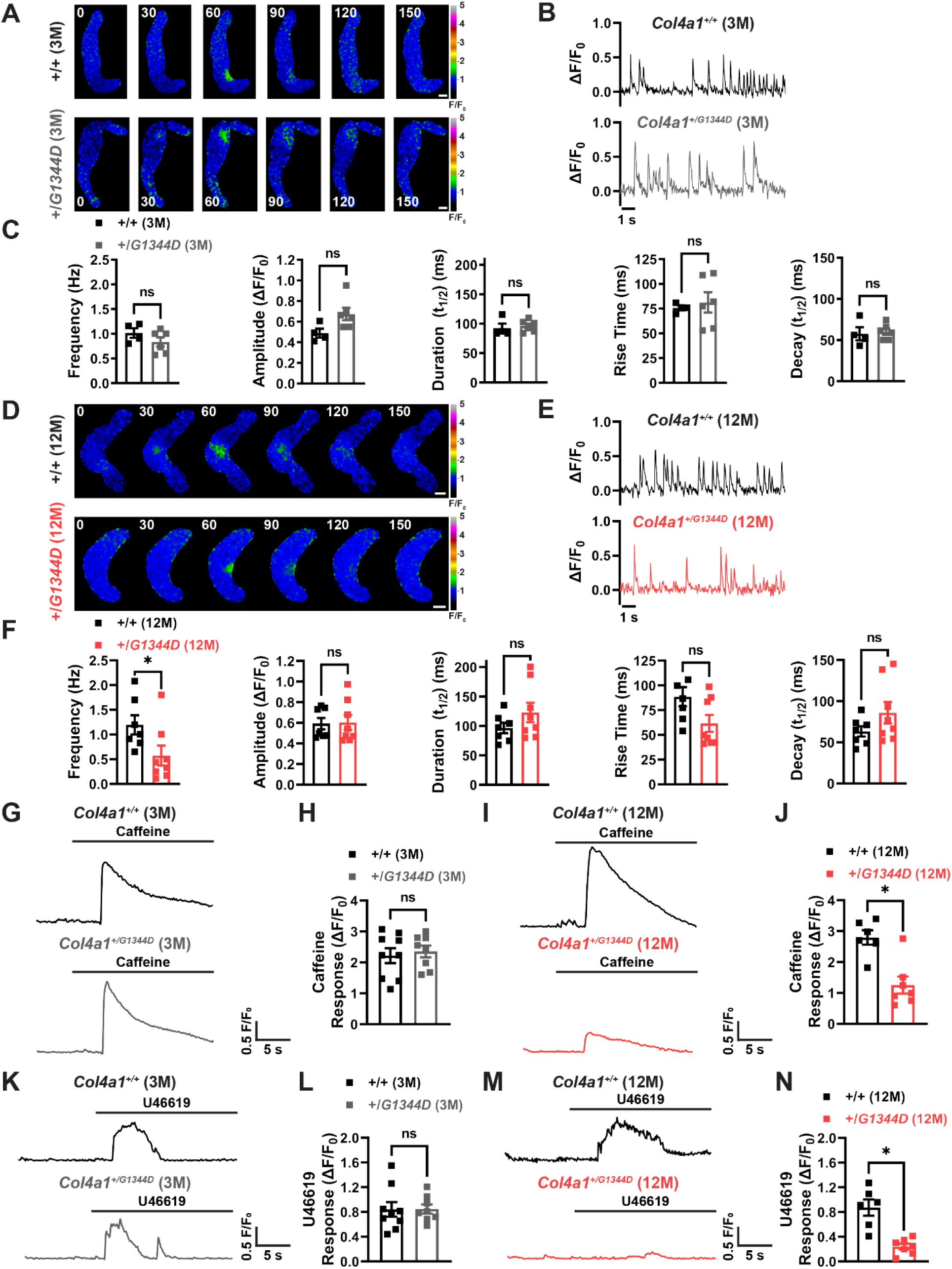
Defective intracellular Ca^2+^ signaling in cerebral artery SMCs from 12 M- old *Col4a1^+/G1344D^* mice. **(A)** Representative time-course spinning disk confocal images exhibiting Ca^2+^ spark events in Fluo-4-AM loaded cerebral artery smooth muscle cells (SMCs) from 3 M-old *Col4a1^+/+^* and *Col4a1^+/G1344D^* mice. Time = milliseconds, scale bar = 5 µm. **(B)** Representative traces of Ca^2+^ spark events in cerebral artery SMCs isolated from 3 M-old *Col4a1^+/+^* and *Col4a1^+/G1344D^* mice presented as changes in fractional fluorescence (ΔF/F_0_) vs. time. **(C)** Summary data showing Ca^2+^ spark frequency, amplitude, duration, rise time, and decay. n = 4-6 cells from 3 animals per group. ns = not significant, unpaired t-test. **(D)** Representative time-course images exhibiting Ca^2+^ spark events in cerebral artery SMCs from 12 M-old *Col4a1^+/+^* and *Col4a1^+/G1344D^*. Time = milliseconds, scale bar = 5 µm. **(E)** Representative traces of Ca^2+^ spark events in cerebral artery SMCs isolated from 12 M-old *Col4a1^+/+^* and *Col4a1^+/G1344D^* mice presented as ΔF/F_0_ vs. time. **(F)** Summary data showing Ca^2+^ spark frequency, amplitude, duration, rise time, and decay. n = 7-8 cells from 3 or 4 animals per group. *P ≤ 0.05, ns = not significant, unpaired t-test. **(G)** Representative traces showing whole-cell F/F_0_ changes in response to caffeine (10 mM) in Fluo-4-AM loaded cerebral artery SMCs from 3 M-old *Col4a1^+/+^* and *Col4a1^+/G1344D^* mice. **(H)** Summary data of the ΔF/F_0_ in response to caffeine. n = 8-9 cells from 3 animals per group. ns = not significant, unpaired t-test. **(I)** Representative traces showing whole-cell F/F_0_ changes in response to caffeine in SMCs from 12 M-old *Col4a1^+/+^* and *Col4a1^+/G1344D^* mice. **(J)** Summary data of the ΔF/F_0_ in response to caffeine. n = 6-7 cells from 3 animals per group. *P ≤ 0.05, unpaired t-test. **(K)** Representative traces showing whole-cell F/F_0_ changes in response to U46619 (100 nM) in Fluo-4-AM loaded cerebral artery SMCs from 3 M-old *Col4a1^+/+^* and *Col4a1^+/G1344D^* mice. **(L)** Summary data of the ΔF/F_0_ in response to U46619. n = 8-9 cells from 3 animals per group. ns = not significant, unpaired t-test. **(M)** Representative traces showing whole-cell F/F_0_ changes in response to U46619 in SMCs from 12 M-old *Col4a1^+/+^* and *Col4a1^+/G1344D^* mice. **(N)** Summary data of the ΔF/F_0_ in response to U46619. n = 6-7 cells from 3 animals per group. *P ≤ 0.05, unpaired t-test.

Fluo-4 AM-loaded SMCs were challenged with the RyR agonist caffeine (10 mM) to investigate how RyR function was affected by the *Col4a1^G1344D^* mutation ^45^. Ca^2+^ signals arising from intracellular release were isolated from Ca^2+^ influx events by acutely removing Ca^2+^ from the extracellular solution immediately before recording. Caffeine-evoked global cytosolic [Ca^2+^] increases did not differ between SMCs from 3 M-old *Col4a1^+/+^* and *Col4a1^+/G1344D^* mice (Figure 4G and H). In contrast, caffeine-induced global Ca^2+^ signals were smaller in amplitude in SMCs from 12 M-old *Col4a1^+/G1344D^* mice than in controls (Figure 4I and J). These data indicate that the release of SR Ca^2+^ from RyRs is impaired in the SMCs of *Col4a1^+/G1344D^* mice in an age-dependent manner.

Fluo-4 AM loaded-SMCs were also treated with the G_q_ protein-coupled thromboxane A2 receptor agonist U46619 (100 nM) to investigate IP_3_R-mediated Ca^2+^ signaling. U46619-induced Ca^2+^ signals did not differ between cerebral artery SMCs from 3 M-old *Col4a1^+/+^* and *Col4a1^+/G1344D^* mice (Figure 4K and L). In contrast, Ca^2+^ signals induced by U46619 were smaller in amplitude in SMCs from 12 M-old *Col4a1^+/G1344D^* mice compared to controls (Figure 4M and N). These data suggest that the release of SR Ca^2+^ from IP_3_Rs is also impaired in the SMCs of *Col4a1^+/G1344D^* mice in an age-dependent manner.

RyRs and IP_3_Rs share a common SR Ca^2+^ pool ^46^. Impairment of Ca^2+^ release through both receptors suggests that the SR [Ca^2+^] of SMCs from 12 M-old *Col4a1^+/G1344D^* mice is lower than that of corresponding controls. Further, diminished SR Ca^2+^ levels likely account for decreased Ca^2+^ spark frequency and diminished STOC and TICC activity.

### 4PBA prevents age-related cerebral artery defects in *Col4a1^+/G1344D^* mice

*Col4a1* mutations impair collagen α1α1α2(IV) secretion *in vitro* and *in vivo* and can be alleviated by treatment with 4PBA ^10,13,26^. The chemical chaperone 4PBA facilitates the trafficking of misfolded mutant proteins and reduces ER/SR stress ^47^. To investigate a potential link between impaired collagen secretion and the cerebral vascular pathology of *Col4a1^+/G1344D^* mice, 4PBA (50 mM) was added to the drinking water of mutant and control animals from birth (postnatal day 0).

We found that 4PBA treatment prevented the diminishment of caffeine-evoked (Figure 5A and B) and U46619-induced (Figure 5C and D) Ca^2+^ signals in cerebral artery SMCs from 12 M-old *Col4a1^+/G1344D^* mice. Ca^2+^ spark frequency was also normalized to control levels in 4PBA-treated *Col4a1^+/G1344D^* mice (Supplemental Figure 3A-C). We also found that 4PBA treatment averted the loss of STOC (Supplemental Figure 3D-E) and TICC (Figure 5E and F) activity. These data demonstrate that age-related defects in intracellular Ca^2+^ signaling that control ion channel activity and membrane potential were prevented by treating *Col4a1^+/G1344D^* mice with 4PBA. 4PBA treatment also prevented loss of myogenic tone (Figure 5G and H) and significantly diminished spontaneous ICH in 12 M-old *Col4a1^+/G1344D^* mice (Figure 5I and J). These data suggest that 4PBA protects against age-dependent defects in myogenic tone and ICH in *Col4a1^+/G1344D^* mice.

**Figure 5.**
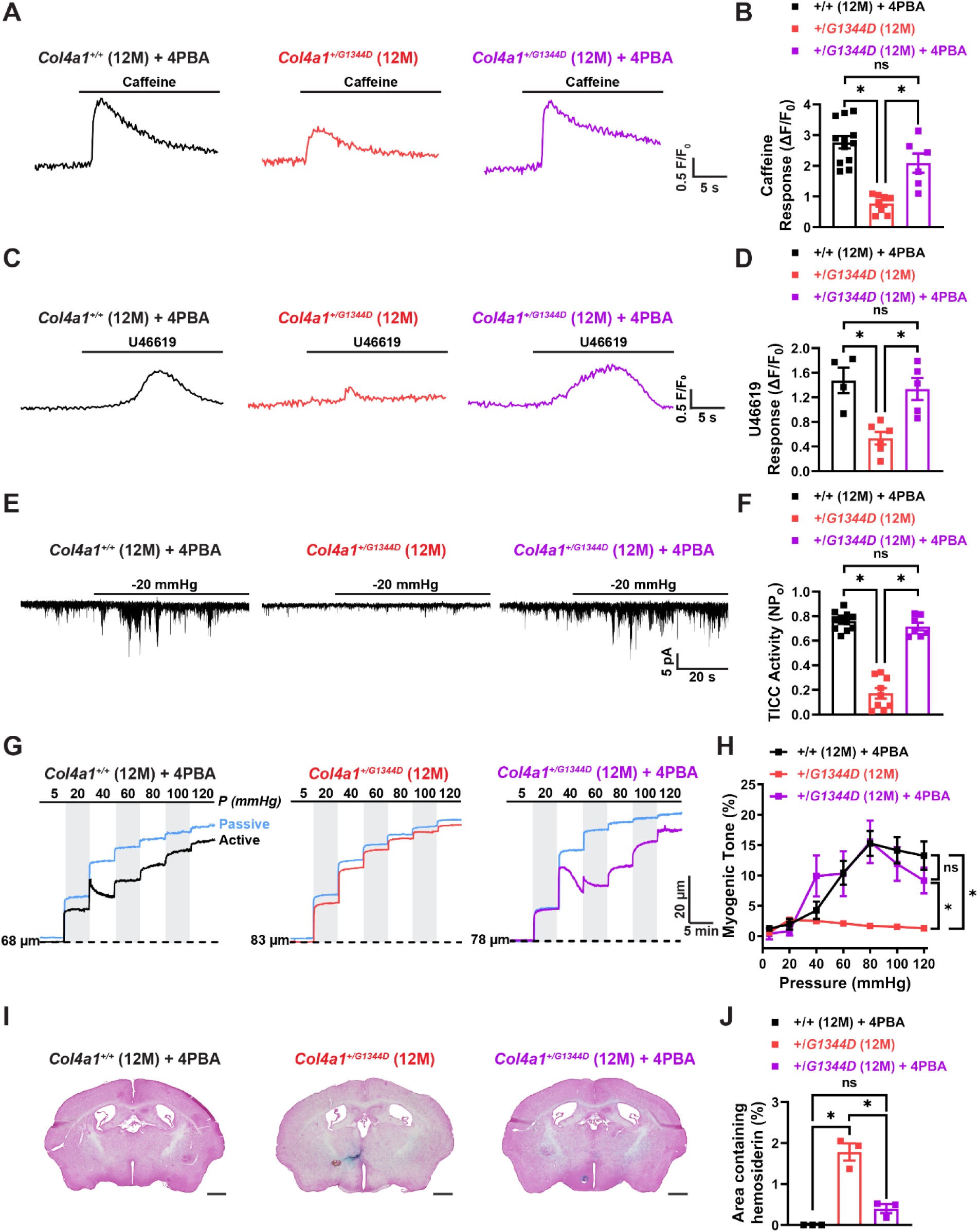
4PBA prevents age-related cerebral artery defects in *Col4a1^+/G1344D^* mice. **(A)** Representative traces showing whole-cell F/F_0_ changes in response to caffeine (10 mM) in Fluo-4-AM loaded cerebral artery SMCs from 4PBA treated 12 M-old *Col4a1^+/+^* and *Col4a1^+/G1344D^* and untreated *Col4a1^+/G1344D^* mice. **(B)** Summary data of the ΔF/F_0_ in response to caffeine. n = 6-12 cells from 3-4 animals per group. *P ≤ 0.05, ns = not significant, one-way ANOVA. **(C)** Representative traces showing whole-cell F/F_0_ changes in response to U46619 (100 nM) in Fluo-4-AM loaded cerebral artery SMCs from 4PBA treated 12 M-old *Col4a1^+/+^* and *Col4a1^+/G1344D^* and untreated *Col4a1^+/G1344D^* mice. **(D)** Summary data of the ΔF/F_0_ in response to U46619. n = 4-6 cells from 3-4 animals per group. *P ≤ 0.05, ns = not significant, one-way ANOVA. **(E)** Representative traces of transient inward cation currents (TICCs) evoked by applying negative pressure (−20 mmHg) to stretch the plasma membrane of voltage-clamped (−70 mV) cerebral artery SMCs isolated from 4PBA treated 12 M-old *Col4a1^+/+^* and *Col4a1^+/G1344D^* and untreated *Col4a1^+/G1344D^* mice. **(F)** Summary data of TICC activity. n = 7-11 cells from 3-4 animals per group. *P ≤ 0.05, ns = not significant, one-way ANOVA. **(G)** Representative recordings of the inner diameter of isolated cerebral arteries from 4PBA-treated 12 M-old *Col4a1^+/+^* and *Col4a1^+/G1344D^* and untreated *Col4a1^+/G1344D^* mice showing the myogenic response to increases in pressure (active) and the dilation of the same arteries when extracellular Ca^2+^ has been removed (passive). **(H)** Summary data of myogenic tone as a function of intraluminal pressure. n = 6 or 9 vessels from 3 or 5 mice. *P ≤ 0.05, ns = not significant, two-way ANOVA. **(I)** Representative images of Prussian blue stained brain sections from 4PBA treated 12 M-old *Col4a1^+/+^* and *Col4a1^+/G1344D^* and untreated *Col4a1^+/G1344D^* mice. Scale bar = 1 mm. **(J)** Summary data presented as percentage of brain area with Prussian blue staining in brain sections (500 µm intervals). N = 3 animals per group. *P ≤ 0.05, ns = not significant, one-way ANOVA.

## Discussion

Despite the enormous impact of the disease, the molecular pathogenesis of cSVD is essentially unknown. Here we utilized the *Col4a1^+/G1344D^* mouse model of Gould syndrome to investigate how this monogenic form of cSVD damages small arteries in the brain during aging. Our findings show that these mice exhibit pathological changes with age, including the loss of myogenic tone, high levels of spontaneous ICH, and brain lesions. Electrophysiology experiments revealed that the loss of myogenic tone is associated with diminished pressure-induced SMC depolarization and decreased activities of BK and TRPM4 channels. In addition, we found that fundamental RyR- and IP_3_R-dependent Ca^2+^ signaling pathways are disrupted in SMCs from 12 M-old *Col4a1^+/G1344D^* mice, likely due to a reduction in SR Ca^2+^ levels. Treating *Col4a1^+/G1344D^* mice with the chemical chaperone, 4PBA negated Ca^2+^ signaling disruptions, increased BK and TRPM4 channel activity, prevented loss of myogenic tone, and reduced spontaneous ICH. We conclude that the age-dependent cSVD pathogenesis of *Col4a1^+/G1344D^* mice results from defects in SMC SR function that lead to reduced Ca^2+^ store load and disrupted intracellular Ca^2+^ signaling. Defects in subcellular Ca^2+^ signals diminish BK and TRPM4 channel activity, disrupting membrane potential regulation and impeding pressure-induced constriction of cerebral arteries. These findings reveal a novel mechanism that contributes to age-dependent cSVD.

Myogenic constriction protects delicate capillaries in the brain from pressure overload during increases in cardiac output and perfusion associated with exercise, stress, and other stimuli ^14^. This vital response is deficient in several monogenetic forms of cSVD ^22,30,48^. However, the defects responsible for the loss of myogenic tone differ between disease models. For example, Dabertrand *et al*. linked the loss of myogenic tone in the CADASIL model with blunted pressure-induced SMC membrane potential depolarization and connected the defect in membrane potential regulation with increased surface expression of K_V_ channels and elevated hyperpolarizing K^+^ current density ^22^. Notably, depletion of the minor membrane phospholipid PIP_2_, leading to the loss of TRPM4 channel activity, is the fundamental defect responsible for the loss of myogenic constriction in a distinct mouse model of Gould syndrome caused by a *Col4a1^G394V^* mutation ^30^. The current study uncovered a distinctive flaw in the SMCs of *Col4a1^+/G1344D^* cSVD mice – impaired Ca^2+^ signaling. Our data show that the release of Ca^2+^ from the SR through RyRs and IP_3_Rs became faulty in *Col4a1^+/G1344D^* mice by 12 M of age. Impaired SR Ca^2+^ release disrupted the activity of BK and TRPM4 channels. BK and TRPM4 channels play opposing roles in regulating SMC membrane potential and contractility. TRPM4 channels depolarize the membrane and promote contractility, whereas BK channels provide negative feedback to limit the magnitude of this response ^49^. When pressure-induced membrane depolarization is impaired, the BK pathway is disengaged and has minimal influence on membrane potential ^50^.

Diminished levels of Ca^2+^ in the SR likely cause Ca^2+^ signaling and ion channel defects in SMCs from 12 M-old *Col4a1^+/G1344D^*. This concept is supported by prior studies showing that a ∼16% decrease in SR [Ca^2+^] diminished STOC frequency by 70% and that a >20% decrease in SR [Ca^2+^] completely abolished IP_3_R-mediated Ca^2+^ release in native SMCs ^51,52^. In many types of cells, store-operated Ca^2+^ entry (SOCE) is protective against decreases in ER/SR [Ca^2+^] ^53^. However, SOCE is virtually undetectable in native, contractile vascular SMCs, promoting their susceptibility to pathogenic SR Ca^2+^ depletion ^50,54,55^. We conclude that SR Ca^2+^ store depletion and the resultant dysregulation of BK, TRPM4, and SMC membrane potential is responsible for the age-related loss of myogenic tone in *Col4a1^+/G1344D^* cSVD mice.

Steady-state free Ca^2+^ levels in the ER/SR are maintained by the release of Ca^2+^ into the cytosol through RyRs and IP_3_Rs, mobile and stationary Ca^2+^-binding proteins in the SR lumen, and transport of Ca^2+^ from the cytosol into the SR by sarco/endoplasmic reticulum ATPase (SERCA) enzymes ^46^. Disruption of one or more of these elements could lead to SR Ca^2+^ defects in *Col4a1^+/G1344D^* mice. Interestingly, *Col4a1^+/G1344D^* mice display impaired secretion and increased intracellular accumulation of collagen α1α1α2(IV), presumably due to difficulties associated with the trafficking of misfolding of mutant proteins out of the ER/SR ^13^. The retention of misfolded proteins causes a pathological condition known as ER/SR stress ^56^ that is associated with the depletion of Ca^2+^ stores ^57^. For example, Yamato *et al.* ^58^ demonstrated that tunicamycin-induced ER stress promotes RyR-mediated Ca^2+^ leak, potentially due to increased reactive oxygen species (ROS) production. Our data showed that treatment with a molecular chaperone prevented SR Ca^2+^ handling defects and loss of myogenic tone and significantly attenuated spontaneous ICH. These data provide evidence that ER/SR stress contributes to SMC Ca^2+^ signaling defects and cSVD pathology of *Col4a1^+/G1344D^* mice. The age dependency of the disease process may reflect the normal functional decline in SR protein homeostasis that is exacerbated by mutant collagen in the SR.

We conclude that age-dependent disruption of Ca^2+^ signaling in SMCs due to SR stress and diminished SR Ca^2+^ store load contributes to cSVD pathology in *Col4a1^+/G1344D^* mice. Future investigations into links between SMC SR stress and Ca^2+^ handling defects in other genetic and sporadic forms of cSVD may provide insight into novel pathogenic mechanisms.

## Materials and Methods

### Chemical and reagents

All chemicals and other reagents were obtained from Sigma-Aldrich, Inc. (St. Louis, MO, USA) unless otherwise specified.

### Animals

Young adult (∼3M) and middle-aged (∼12M) male and female littermate *Col4a1^+/+^* and *Col4a1^+/G1344D^* mice were used in this study. The *Col4a1^G1344D^* mutation was backcrossed to C57BL/6J mice for over 20 generations ^59^. Animals were maintained in individually ventilated cages (≤5 mice/cage) with *ad libitum* access to food and water in a room with controlled 12-hour light and dark cycles. All animal care procedures and experimental protocols involving animals complied with the National Institutes of Health (NIH) *Guide for the Care and Use of Laboratory Animals* and were approved by the Institutional Animal Care and Use Committees at the University of Nevada, Reno, and University of California, San Francisco.

### 4PBA treatment

Mice were provided with 4-phenylbutyrate (4PBA; 50 mM; Scandinavian Formulas Inc., Sellersville, PA, USA) from birth in drinking water that was refreshed weekly as described previously ^26^.

### Histological analysis

Isoflurane anesthetized mice were transcardially perfused with ice-cold phosphate-buffered saline (PBS) followed by ice-cold 4% paraformaldehyde (Fisher, Waltham, MA, USA) in PBS. Brains were postfixed with 4% paraformaldehyde for 24 hours at 4°C, cryoprotected in 30% sucrose in PBS, and embedded in optimal cutting temperature compound (Fisher). Coronal cryosections (35 µm) regularly spaced (500 µm) along the rostrocaudal axis were stained with a Prussian blue and nuclear fast red stain kit (Abcam, Cambridge, UK) following the manufacturer’s protocol. Images were acquired with a BZ-X700 microscope using BZ-X Viewer 1.3.0.5 software and stitched with BZ-X Analyzer 1.3.0.3 software (Keyence, Osaka, Japan). The percentage of brain area with Prussian blue staining for each section was calculated with ImageJ software (v2.3.0/1.53f; NIH, Bethesda, MD, USA). Hemorrhage severity was expressed as the average percentage of hemosiderin surface area on ∼21 sections for each brain.

### *In vivo* magnetic resonance imaging

All *in vivo* magnetic resonance (MR) experiments were conducted on a 14.1 Tesla vertical MR system (Agilent Technologies, Palo-Alto, CA) equipped with 100 G/cm gradients and a single tuned millipede ^1^H proton coil (Ø_I_ = 40 mm). For each imaging session, mice were anesthetized using isoflurane (1-1.5% in O_2_), positioned in a dedicated cradle maintaining constant anesthesia, and placed in the MR bore; respiration and temperature were continuously monitored during all acquisitions to ensure animal well-being and data reproducibility. Susceptibility weighted imaging (SWI) was performed to acquire high-resolution images of the mouse brain using the following parameters: gradient-echo scheme, field of view (FOV) = 20x 20 mm^2^, matrix = 256 x 256, 16 slices, 0.4 mm slice thickness, 0.1 mm interslice gap, number of averages = 16, echo time (TE)/repetition time (TR) = 4.60/140ms, flip angle = 10 degrees. T2-weighted (T2W) images were also acquired in a fast-spin-echo scheme to evaluate anatomical brain structures, using the same FOV geometry as SWI and the following parameters: number of averages = 8, TE/TR = 21.38/2500ms, flip angle = 90 degrees. The fast-spin-echo based T2W imaging is less sensitive to distortion artifacts caused by B_0_ inhomogeneity than SWI and is thus used for the atlas-based brain structure registration as described below.

Using in-house software, hypointense lesions corresponding to ICH were quantified based on manual segmentation of the SWI images. To investigate the number and distribution of hypointense SWI lesions across the brain, an open-sourced atlas-based imaging data analysis pipeline (AIDAmri) ^60^ was customized to register T2W brain images to the Allen Brain Reference Atlas ^61^. Based on the registration result, the normalized lesion volumes for each brain region were summed up across all animals, and an intra-group region-wise volumetric analysis was performed.

### Isolation of cerebral arteries and SMCs

Mice were euthanized by decapitation under isoflurane anesthesia. Cerebral arteries were carefully isolated from the brain in Ca^2+^-free Mg^2+^ physiological saline solution (Mg^2+^-PSS) consisting of 140 mM NaCl, 5 mM KCl, 2 mM MgCl_2_, 10 mM HEPES, and 10 mM glucose (pH 7.4, adjusted with NaOH), with 0.5% bovine serum albumin (BSA). Native SMCs for patch-clamp experiments were obtained by initially digesting isolated arteries in 1 mg/mL papain (Worthington Biochemical Corporation, Lakewood, NJ, USA), 1 mg/mL dithiothreitol (DTT), and 10 mg/mL BSA in Ca^2+^-free PSS at 37°C for 12 minutes, followed by a 14-minute incubation with 1 mg/mL type II collagenase (Worthington Biochemical Corporation). A single-cell suspension was prepared by washing digested arteries three times with Mg^2+^-PSS and triturating with a fire-polished glass pipette. All cells used for this study were freshly dissociated on the day of experimentation.

### Pressure myography

The current best practice guidelines for pressure myography experiments were followed ^32^. Arteries were mounted between two glass cannulas (outer diameter, 40–50 μm) in a pressure myograph chamber (Living Systems Instrumentation, St Albans City, VT, USA) and secured by a nylon thread. Intraluminal pressure was controlled using a servo-controlled peristaltic pump (Living Systems Instrumentation), and preparations were visualized with an inverted microscope (Accu-Scope Inc., Commack, NY, USA) coupled to a USB camera (The Imaging Source LLC, Charlotte, NC, USA). Changes in luminal diameter were assessed using IonWizard software (version 7.2.7.138; IonOptix LLC, Westwood, MA, USA). Arteries were bathed in warmed (37°C), oxygenated (21% O_2_, 6% CO_2_, 73% N_2_) PSS (119 mM NaCl, 4.7 mM KCl, 21 mM NaHCO_3_, 1.17 mM MgSO_4_, 1.8 mM CaCl_2_, 1.18 mM KH_2_PO_4_, 5 mM glucose, 0.03 mM EDTA) at an intraluminal pressure of 5 mmHg. Following equilibration for 15 min, intraluminal pressure was increased to 110 mmHg, and vessels were stretched to their approximate *in vivo* length, after which pressure was reduced back to 5 mmHg for an additional 15 min. Vessel viability was assessed for each preparation by evaluating vasoconstriction response to high extracellular [K^+^] PSS, made isotonic by adjusting the [NaCl] (60 mM KCl, 63.7 mM NaCl). Arteries that showed less than 10% constriction in response to elevated [K^+^] were excluded from further investigation.

Myogenic tone was assessed by raising the intraluminal pressure stepwise from 5 mmHg to 120 mmHg in 20 mmHg increments (5 to 20 mmHg for the first step). The active diameter was obtained by allowing vessels to equilibrate for at least 5 minutes at each pressure or until a steady-state diameter was reached. Following completion of the pressure-response study, intraluminal pressure was lowered to 5 mmHg, and arteries were superfused with Ca^2+^-free PSS supplemented with EGTA (2 mM) and the voltage-dependent Ca^2+^ channel blocker diltiazem (10 μM) to inhibit SMC contraction, after which passive diameter was obtained by repeating the stepwise increase in intraluminal pressure. Myogenic tone at each pressure step was calculated as myogenic tone (%) = [1 – (active lumen diameter/passive lumen diameter)] × 100.

### Membrane potential

For measurements of SMC membrane potential, cerebral arteries were isolated, cannulated, and viability confirmed as described above. SMC membrane potential was recorded from intact pressurized arteries at intraluminal pressures of 20 mmHg and 80 mmHg. SMCs were impaled through the adventitia with glass intracellular microelectrodes (100-200 MΩ) backfilled with 3M KCl. Membrane potential was recorded using an Electro 705 amplifier (World Precision Instruments, Sarasota, FL, USA). Analog output from the amplifier was recorded using Ionwizard software (version 7.5.1.162; IonOptix). Criteria for acceptance of membrane potential recordings were 1) an abrupt negative deflection of potential as the microelectrode was advanced into a cell, 2) stable membrane potential for at least 30 seconds, and 3) an abrupt change in the potential to ∼0 mV after the electrode was retracted from the cell.

### Electrophysiological recordings

Enzymatically isolated native SMCs were transferred to a recording chamber (Warner Instruments, Holliston, MA, USA) and allowed to adhere to glass coverslips for 20 minutes at room temperature. Recording electrodes (3–5 MΩ) were pulled on a model P-87 micropipette puller (Sutter Instruments, Novato, CA, USA) and polished using a MF-830 MicroForge (Narishige Scientific Instruments Laboratories, Long Island, NY, USA). Currents were recorded at room temperature using an Axopatch 200B amplifier (Molecular Devices, Sunnyvale, CA, USA) equipped with an Axon CV 203BU headstage and Digidata 1440A digitizer (Molecular Devices) for all patch-clamp electrophysiology experiments. Clampex and Clampfit (version 10.2; Molecular Devices) were used for data acquisition and analysis.

The bathing solution composition for perforated-patch recordings of Kv currents, STOCs, and TICCs was 134 mM NaCl, 6 mM KCl, 1 mM MgCl_2_, 2 mM CaCl_2_, 10 mM HEPES, and 10 mM glucose (pH 7.4, adjusted with NaOH). The patch pipette solution contained 110 mM K-aspartate, 1 mM MgCl_2_, 30 mM KCl, 10 mM NaCl, 10 mM HEPES, and 5 μM EGTA (pH 7.2, adjusted with KOH). Amphotericin B (200 µM) was included in the pipette solution to gain electrical access. Whole-cell K^+^ currents were recorded using a step protocol (−60 to +60 mV in 10 mV, 250 ms steps) from a holding potential of −80 mV. The BK channel blocker paxilline (1 µM) was included in the bath solution when Kv currents were recorded. Whole-cell K^+^ currents were recorded in the absence and presence of K_V_ channel blocker 4-aminopyridine (5 mM), and the Kv component was determined by subtraction. Summary current-voltage (I-V) plots were generated using values obtained from the last 50 ms of each step. STOCs were recorded from SMCs voltage-clamped over a range of membrane potentials (−60 to 0 mV; 10 mV steps). TICCs were recorded from SMCs voltage-clamped at −70 mV; membrane stretch was delivered by applying negative pressure through the recording electrode using a pressure clamp system (HSPC-1; ALA Scientific Instruments Inc., Farmingdale, NY, USA). TICC activity was calculated as the sum of the open channel probability (NP_o_) of multiple open states of 1.75 pA ^62^.

Whole-cell TRPM4 currents were recorded in a bath solution consisting of 156 mM NaCl, 1.5 mM CaCl_2_, 10 mM glucose, 10 mM HEPES, and 10 mM TEACl (pH 7.4, adjusted with NaOH). The patch pipette solution contained 156 mM CsCl, 8 mM NaCl, 1 mM MgCl_2_, 10 mM HEPES (pH 7.4, adjusted with NaOH), and a free Ca^2+^ concentration of 200 µM, adjusted using the appropriate amount of CaCl_2_ and EGTA, as determined with Max-Chelator software (WEBMAXC standard, available at https://somapp.ucdmc.ucdavis.edu/pharmacology/bers/maxchelator/webmaxc/webmaxcS.htm. Whole-cell cation currents were evoked by applying 400 ms voltage ramps from −100 to +100 mV from a holding potential of −60 mV. Voltage ramps were repeated every 2 s for 300 s. The selective TRPM4 inhibitor 9-phenanthrol (30 μM) was applied after peak TRPM4 current was recorded (∼100 s). Whole-cell TRPM4 current amplitude was expressed as the 9-phenanthrol-sensitive current at +100 mV.

### Quantitative droplet digital PCR

Total RNA was extracted from isolated cerebral arteries by homogenization in TRIzol reagent (Invitrogen, Waltham, MA, USA), followed by purification using a Direct-zol RNA microprep kit (Zymo Research, Irvine, CA, USA) with on-column DNAse treatment. RNA concentration was determined using an RNA 6000 Pico Kit run on a Bioanalyzer 2100 running Agilent 2100 Expert Software (B.02.11; Agilent Technologies, Santa Clara, CA, USA). RNA was converted to cDNA using iScript cDNA Supermix (Bio-Rad, Hercules, CA, USA). Quantitative droplet digital PCR (ddPCR) was performed using QX200 ddPCR EvaGreen Supermix (Bio-Rad), custom-designed primers (Supplementary Table 1), and cDNA templates. Generated droplet emulsions were amplified using a C100 Touch Thermal Cycler (Bio-Rad), and the fluorescence intensities of individual droplets were measured using a QX200 Droplet Reader (Bio-Rad) running QuantaSoft (version 1.7.4; Bio-Rad). Analysis was performed using QuantaSoft Analysis Pro (version 1.0596; Bio-Rad).

### Ca^2+^ imaging

Images were acquired using an iXon 897 EMCCD camera (Andor; 16 × 16 µm pixel size) coupled to a spinning-disk confocal head (CSU-X1; Yokogawa) with a 100× oil-immersion objective (Olympus; NA 1.45) at an acquisition rate of 33 frames per second (fps). Custom software (SparkAn; https://github.com/vesselman/SparkAn) provided by Dr. Adrian D. Bonev (University of Vermont) was used to analyze the properties of Ca^2+^ sparks and whole-cell RyR and IP_3_R Ca^2+^ signals.

To image Ca^2+^ sparks, a suspension of freshly isolated SMCs was placed in a glass-bottom 35 mm dish, and cells were allowed to adhere to the glass coverslip for 20 min at room temperature. SMCs were then loaded with the Ca^2+^-sensitive fluorophore, Fluo-4-AM (1 µM; Invitrogen), in the dark for 20 minutes at room temperature in Mg^2+^-PSS. Cells were subsequently washed three times with Ca^2+^-containing PSS and incubated at room temperature for 20 min in the dark to allow sufficient time for Fluo-4 de-esterification. The threshold for Ca^2+^ spark detection was defined as local increases in fluorescence ≥0.2 ΔF/F_0_.

To image RyR and IP_3_R Ca^2+^ signals, SMCs were plated and loaded with Fluo-4-AM as stated above, but immediately before imaging, cells were washed three times and imaged in Mg^2+^-PSS to eliminate signals resulting from the influx of Ca^2+^. Caffeine (10 mM) or U46619 (100 nM; Enzo Biochem, Farmingdale, NY) were applied to the bath.

### Statistical analysis

All summary data are presented as means ± SEM. Statistical analyses were performed, and graphs were constructed using Prism version 9.3.1 (GraphPad Software, San Diego, CA, USA). The value of n refers to the number of cells for patch-clamp electrophysiology and Ca^2+^ imaging experiments, the number of arteries for pressure myography and membrane potential experiments, and the number of animals for histological analyses, magnetic resonance imaging, and ddPCR experiments. Statistical analyses were performed using unpaired Student’s t-tests and one-way and two-way analysis of variance (ANOVA). A P value < 0.05 was considered statistically significant for all analyses.

### Grant support

This study was supported by grants from the National Institutes of Health (NHLBI R35HL155008, R01HL091905, R01HL137852, R01HL139585, R01146054, and R01HL122770, NIGMS P20GM130459 to S.E., and NINDS RF1NS110044 and R33NS115132 to M.M.C, D.B.G, and S.E.; The Transgenic Genotyping and Phenotyping Core at the COBRE Center for Molecular and Cellular Signaling in the Cardiovascular System, University of Nevada, Reno, is maintained by a grant from NIH/NIGMS (P20GM130459 Sub#5451), as is the High Spatial and Temporal Resolution Imaging Core at the COBRE Center for Molecular and Cellular Signaling in the Cardiovascular System, University of Nevada, Reno (P20GM130459 Sub#5452). The UCSF Department of Ophthalmology is supported by P30EY002162 and an unrestricted grant from Research to Prevent Blindness. This study was also supported by a grant from the American Heart Association (23CDA1054627 to P.T.).

### Author contributions

S.E. and D.B.G. initiated and supervised the project. S.E. designed the experiments. E.Y. performed brain histology, pressure myography, sharp electrode and patch-clamp electrophysiology, Ca^2+^ imaging, and ddPCR experiments. X.W. and X.G. performed magnetic resonance imaging experiments. S.A. and A.S.S. performed patch-clamp electrophysiology experiments. P.T. performed pressure myography experiments. C.L.D. performed animal colony management. E.Y., S.A., A.S.S., P.T., X.G., and S.E. analyzed the data. E.Y. and S.E. drafted the manuscript and prepared the figures. E.Y., C.L.D, M.M.C., D.B.G., and S.E. revised the manuscript.

### Data and materials availability

All data needed to evaluate the conclusions in the paper are present in the paper or the Supplementary Materials

