## Supplemental Materials for "Impaired intracellular Ca^2+^ signaling contributes to age-related cerebral small vessel disease in *Col4a1* mutant mice"

**This PDF file includes:**

- Supplemental Figures 1-3
- Supplemental Table 1

### Supplemental Figures

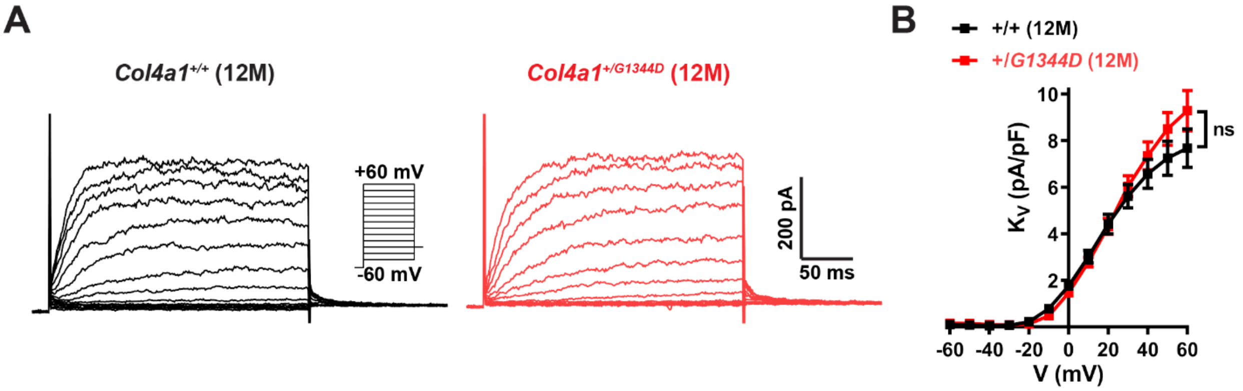


**Supplemental Figure 1 – SMC Kv currents do not differ between 12 M-old *Col4a1*$^{+/+}$ and *Col4a1*$^{+/G1344D}$ mice. (A)** Representative recording of 4-aminopyridine (5 mM)-sensitive Kv currents in freshly isolated cerebral artery smooth muscle cells from 12 M-old *Col4a1*$^{+/+}$ and *Col4a1*$^{+/G1344D}$ mice. Currents were elicited by applying voltage pulses (250 ms) from -60 to +60 mV in 10 mV steps with BK channel blocker paxilline (1 µM). **(B)** Summary data of Kv current at each command potential, normalized to cell capacitance. n = 10-11 cells from 3 animals per group. ns = not significant, two-way ANOVA.

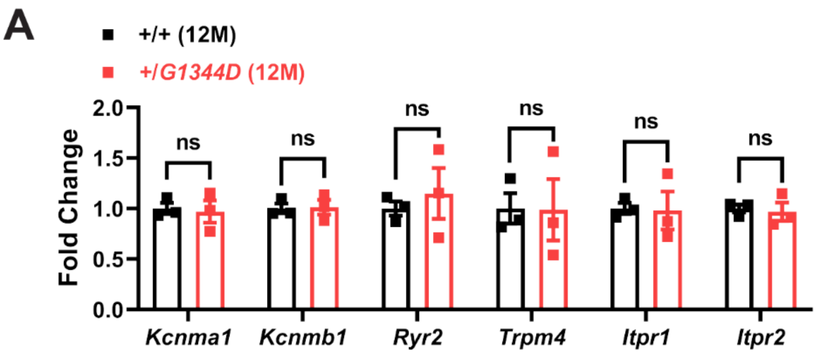


**Supplemental Figure 2 – mRNA expression of BK and TRPM4 signaling pathway components.** mRNA transcript levels of genes involved in the spontaneous transient outward current (*Kcnma1, Kcnmb1, Ryr2)* and transient inward cation current (*Trpm4, Itpr1, Itpr*2) pathways in isolated cerebral arteries from 12 M-old *Col4a1*$^{+/+}$ and *Col4a1*$^{+/G1344D}$ mice measured by ddPCR and presented as fold change over control mice. n = 3 animals per group. ns = not significant, unpaired t-test.

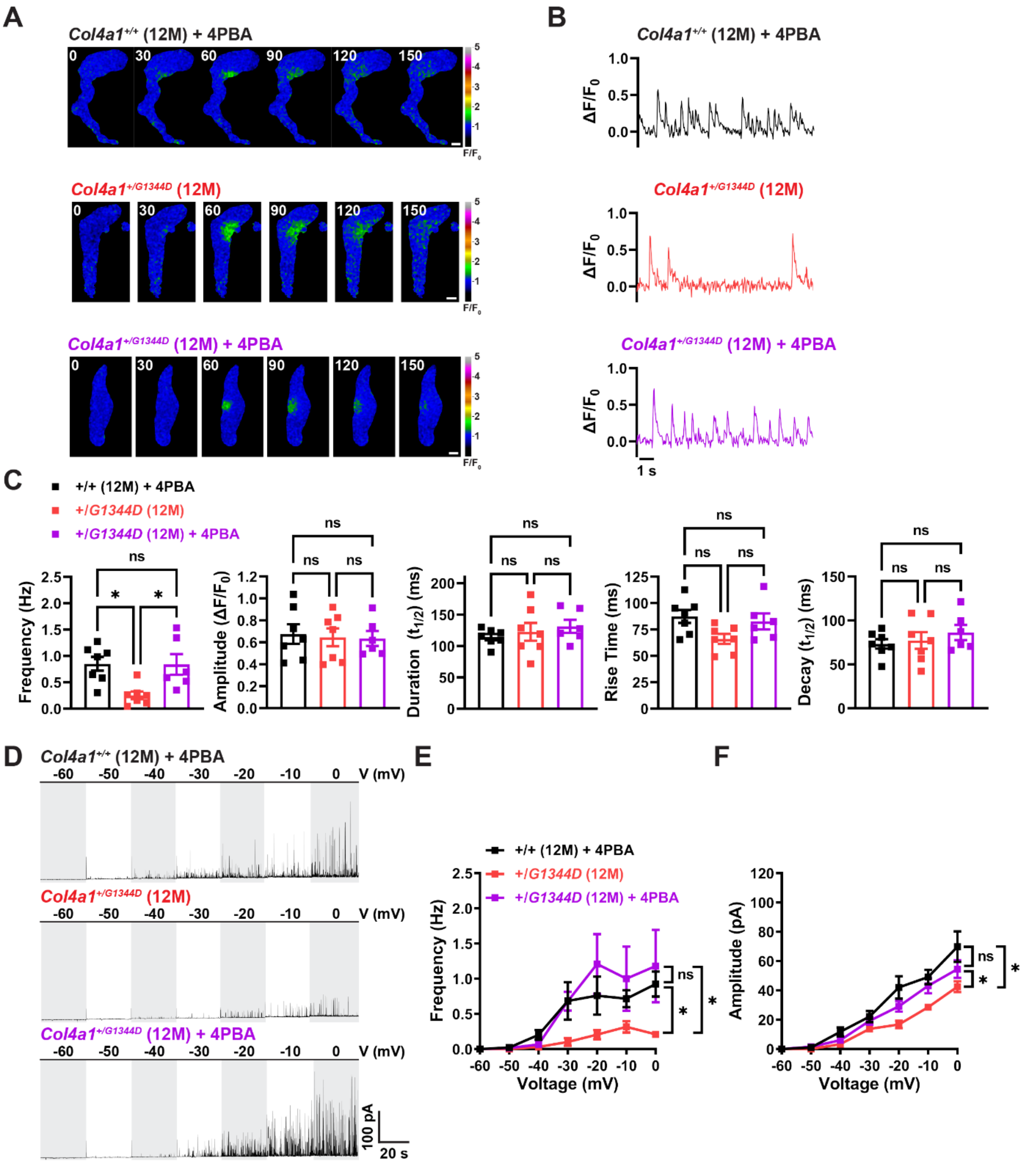


**Supplemental Figure 3 - 4PBA prevents age-dependent decreases in $Ca^{2+}$ spark frequency and BK channel activity. (A)** Representative time-course spinning disk confocal images exhibiting $Ca^{2+}$ spark events in Fluo-4-AM loaded cerebral artery smooth muscle cells (SMCs) from 12 M-old 4PBA treated *Col4a1*$^{+/+}$ and *Col4a1*$^{+/G1344D}$

mice and untreated *Col4a1*$^{+/G1344D}$ mice. Time = milliseconds, scale bar = 5 μm. **(B)** Representative traces of $Ca^{2+}$ spark events presented as changes in fractional fluorescence ($\Delta F/F_0$) vs. time. **(C)** Summary data showing $Ca^{2+}$ spark frequency, amplitude, duration, rise time, and decay. n = 6-7 cells from 3-7 animals per group. *P ≤ 0.05, ns = not significant, one-way ANOVA. **(D)** Representative traces of spontaneous transient outward currents (STOCs) in freshly isolated cerebral artery SMCs from 12 M-old 4PBA treated *Col4a1*$^{+/+}$ and *Col4a1*$^{+/G1344D}$ mice and untreated *Col4a1*$^{+/G1344D}$ mice over a range of membrane potentials (-60 to 0 mV). Summary data showing STOC frequency **(E)** and amplitude **(F)** at each command potential. n = 8-11 cells from 3-6 animals per group. *P ≤ 0.05, ns = not significant, two-way ANOVA.

**Supplemental Table 1. Primers used for quantitative ddPCR.**

| **Gene** | **Forward primer (5'-3')** | **Reverse primer (5'-3')** |
|---|---|---|
| *Kcnma1* | GCTTAAGCTCCTGATGATAGCC | AAGGTGGTTCCCAGGGTTAA |
| *Kcnmb1* | ATGGGCCATGCTGTATCACA | TGTCCAGGTTCCTGGGGATATA |
| *Ryr2* | TGGAGGACATGCATCCAACA | TCCTATGCCTGACAAGAACTCC |
| *Trpm4* | AGGGCTCTTGTGAAAGCCTG | TCCCCACGGAAAAGTTCAC |
| *Itpr1* | AACGTGGGCCACAACATCTA | CCAGGTTTCAGCATGGTTTGAA |
| *Itpr2* | CCTCAAGACAACCTGCTTCA | TGATGTGCTCCTCAAAGGAC |
